# The individual and combined benefits of different non-equilibrium proofreading mechanisms

**DOI:** 10.1101/2022.07.05.498791

**Authors:** Adélaïde A. Mohr, Daniel M. Busiello, Stefano Zamuner, Paolo De Los Rios

## Abstract

Genome duplication, transcription and translation are among many crucial cellular processes that need to be performed with high fidelity. However those extremely low error rates cannot be explained with simple equilibrium thermodynamic considerations. They instead require considering irreversible, energy consuming reactions in the overall mechanism. We develop here a model of substrates selection comprising energy consuming steps and which aims at selecting right substrates among wrong ones. With this model, we investigate the impact of energy consumption on the accuracy and the speed of the selection, as well as different selection strategies. The model presented here encompasses the classic kinetic proofreading scheme and a different mechanism whereby the rates of the energy consuming step are modulated by the nature of the substrate. We show that, in our framework, the fastest and most accurate selection strategy relies on a combination of both mechanisms. A structurally and biochemically informed coarse-grained description of real biological processes such as DNA replication and protein translation, traditionally used as examples of kinetic proofreading at work, shows that, as a matter of fact, a combination of both mechanisms explored here is exploited.

## Introduction

The ability to discriminate between correct and wrong substrates is essential for many biological processes, and in particular when information is transferred during transcription and translation. Indeed, from DNA replication to protein synthesis, the ability to read information and select the correct nucleotide or anti codon is crucial for the survival of all living organisms and of their progeny. From a thermodynamic perspective, high fidelity is difficult to achieve because the free-energy differences Δ*G* = *G_R_* – *G_W_* that should allow choosing the right (*R*) rather than the wrong (*W*) substrate are often in the range of only a few *k_B_T*. Therefore, it becomes impossible for the error rate of these processes to be lower than one wrong substrate in a few hundreds, according to ratio of their Boltzmann probabilities, exp(–Δ*G/k_B_T*). Yet, the error rate in translation has been measured to be as low as ~ 10^-4^ [1], and a staggering ~ 10^-10^ for DNA replication [2]. To overcome this conundrum, it has been proposed, consistent with biochemical observations, that some irreversible, energy-consuming steps must be integral parts of these highly accurate processes, thus unshackling them from the bounds of equilibrium thermodynamics.

Hopfield and Ninio [3, 4] independently proposed, almost five decades ago, a proofreading scheme able to reproduce these exceptionally low error rates. By repeating the selection through *n* independent steps, separated by irreversible reactions, the error rate can be reduced to (exp(–Δ*G/k_B_T*))^*n*^. Several subsequent studies have chosen similar approaches to investigate different selection methods [5], more complex kinetic pathways [6] or different characteristics of these proofreading schemes [7, 8, 9, 10].

Although many of these studies acknowledge the need for energy dissipation, possibly in the form of nucleotide (ATP or GTP, henceforth NTP for generality) consumption for proper functioning, few of them explicitly respect simple thermodynamics constraints, such as detailed balance in equilibrium conditions.

In this work, we propose a general model of proofreading that explicitly considers the energy available to the system as a function of the NTP and NDP concentration (NDP is the product of the hydrolysis of NTP). This framework encompasses the classical kinetic proofreading scheme as well as other, not mutually exclusive, means of selection. We compare the behaviour of these selection methods in different energy conditions, discuss their limitations and advantages. We find that Hopfield’s kinetic proofreading scheme is greatly limited by required slow transitions implying a strong trade-off between the accuracy and the speed of the selection. However, its combination with another selection mechanism allows to lift that limitation and leads to an improvement of both the accuracy and the velocity. Finally, the study of two different real biological systems highlights the use of similar combinations of selection methods.

## Results

### Proofreading model

A model capturing the basic proofreading scheme, while allowing the correct accounting of the energy that is available to the process, is represented in Fig. 1. The enzyme responsible for the substrate selection and product formation can either be in an NTP bound-state, *E*, or in an NDP bound-state *E*\*. It original state *E* can either bind to the right, *R*, or wrong, *W*, substrate. After substrate binding, the enzyme can evolve into an intermediate state, *E*S* (*S* is either *R* or *W*) that is reached either through an energy-consuming reaction (here hydrolysis of NTP into NDP; blue arrows in the figure) or by nucleotide exchange (release of NTP and binding of NDP marked by the black transition lines between *E* and *E\** states). If the substrate is released (*E*S* → *E*\*), the enzyme can then exchange NDP for NTP. Otherwise the substrate is turned into a product (*E*S* → *E*P_S_*), which can then unbind before or after NDP is exchanged for NTP.

**Figure 1:**
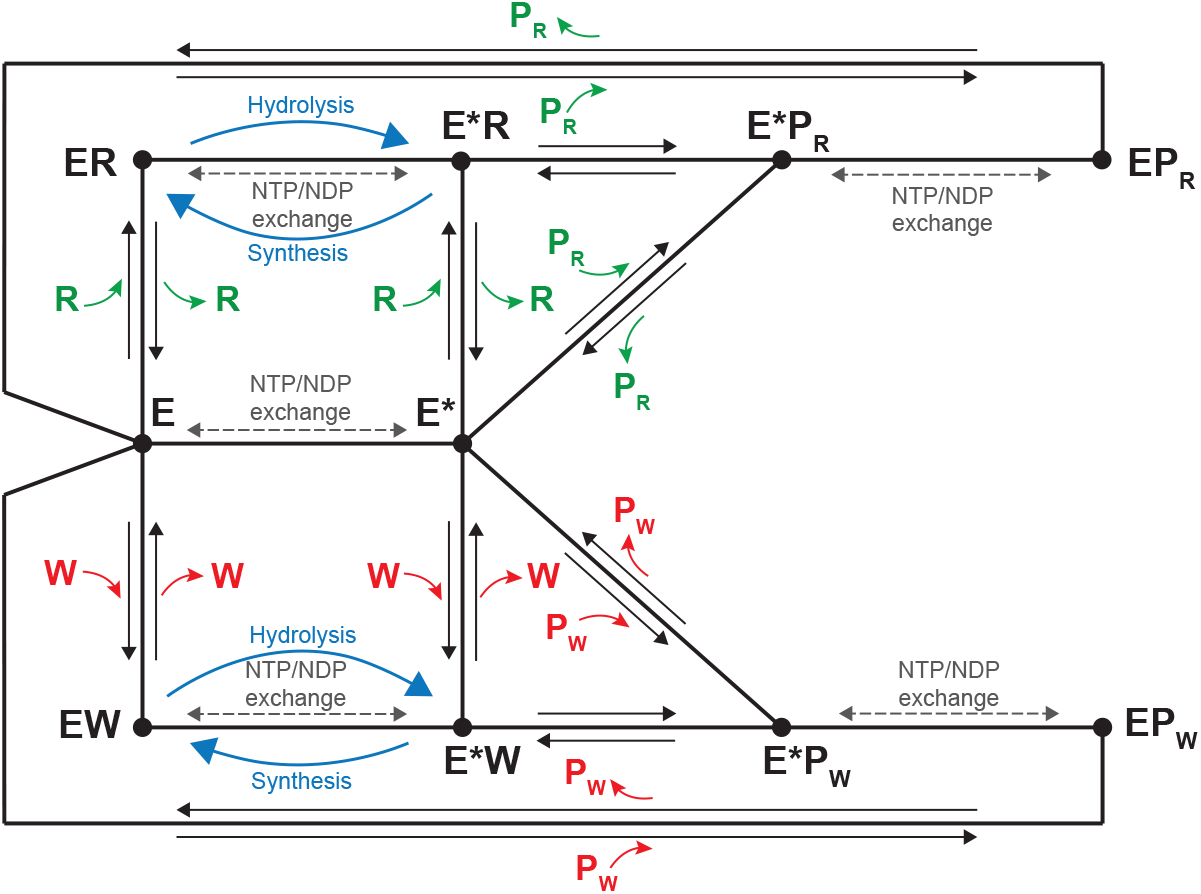
Thermodynamically consistent proofreading scheme. *E* is the enzyme in its NTP-state and *E\** the enzyme in its NDP-state, *R* and *W* correspond to the right and wrong substrate and *P_R_* and *P_W_* the products associated to the respective substrate. Transition between NTP and NDP-states enzymes can either happen through nucleotide exchange (gray dashed arrows) or via NTP hydrolysis or NDP synthesis (blue arrows) when the enzyme is bound to the substrate (*R* or *W*). This is due to the approximation of very slow hydrolysis and synthesis, negligible compared to exchange, unless stimulated by a substrate. Other transitions in the scheme simply corresponds to binding or unbinding to substrates or products.

In this model, we assume the hydrolysis and synthesis to be very slow processes which can be neglected compared to the nucleotide exchange. Only the presence of a substrate, right or wrong, is able to stimulate enough these reactions, explaining why those transitions are only present between *ES* and *E*S*.

A centerpiece of this model is the reversibility of all reactions, which allows deriving a correct relation between the ability of the system to discriminate right substrates from wrong ones and the available energy. Irreversibility or, to be more correct, directionality of reactions emerges from the unbalance between hydrolysis/synthesis and nucleotide exchange. This unbalance is tuned by the parameter *α* = [NTP]/[NDP] representing the amount of energy available to the system. At equilibrium *α* = *α*_eq_ = 10^-6^, corresponding to the equilibrium ratio between NTP and NDP. For any value of *α* > *α*_eq_, the system is pushed out of equilibrium, breaking the detailed balance and leading to directionality in the system reactions. In this work, we usually compare the equilibrium case, *α* = *α*_eq_, with the physiological one, *α* =10 [11].

In what follows, we consider a fixed concentration of substrate and products. In particular, [*P_W_*] = [*P_R_*] = 0 as we assume that products are immediately extracted from the solution. As a consequence, there is a constant net flux flowing through the network even at equilibrium due to these chemostatted concentrations.

Both at equilibrium and out of equilibrium, the system reaches a steady-state. In this state, we compute the flux of wrong and correct products, *J_w_* and *J_r_*, released by the enzyme. To measure the fidelity of the process, we define, as in [3], the error rate 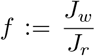. The quantities *J_r_* and *f* will be used to assess the speed and accuracy of different selection strategies. Further mathematical details on the model are given in the appendices.

### Kinetic proofreading: Trade-off between speed and accuracy

The original Hopfield-Ninio selection scheme, represented in Fig. 2, is called **kinetic proofreading** and is based on the presence of the two discriminating steps separated by an irreversible reaction. The initial discriminating step acts as the usual selection step of an enzyme in the classic Michaelis-Menten description, while the second discriminating step allows to reject wrong substrates, acting thus as a proofreading mechanism. The kinetic proofreading scheme can be obtained from the model introduced above by considering that the synthesis and exchange reactions between *ES* to *E*S* are infinitely slow, while the exchange reaction between *E*\* and *E* is infinitely fast, and *α* → ∞. Whereas at equilibrium (*i.e.* no irreversible step), the error rate would have a simple linear dependence on the ratio of the dissociation constants, the presence of an irreversible step allows the system to probe the substrates twice, resulting in

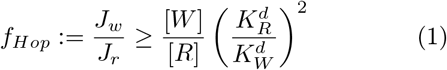

a relation that can be simply derived within a Michaelis-Menten approach.

**Figure 2:**
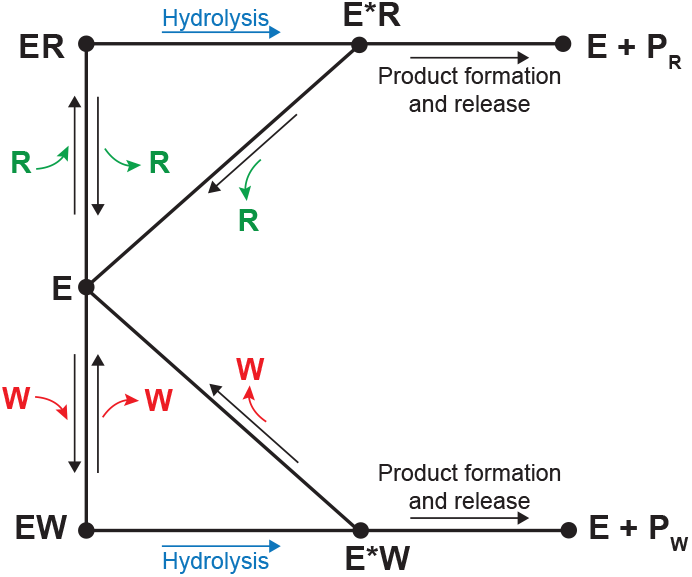
Hopfield’s proofreading scheme. Under Hopfield’s hypothesis the system is symmetric in both the hydrolysis and product formation rates and those rates are slow. It is worth noting the unidirectionality of the transitions in hydrolysis, product formation and release and the release of the substrate in state *E*S*

Since the Hopfield model represented in Fig.2 is a special case of the one presented in this work (Fig. 1), its outcome can be recovered in a suitable limit, with the advantage that, being it thermodynamically consistent, the interplay between energy consumption, fidelity and velocity can be fully appreciated.

To do so, we assume our system to be completely symmetric between the correct and incorrect substrates except for the unbinding rates: 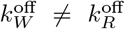 and 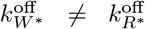. In agreement with Hopfield’s assumption, we set these unbinding rates as large as 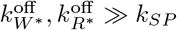, where *k_SP_* is the product production rate for both correct and wrong substrates.

In Fig.3A we show the behavior of the error rate *f* as a function of the hydrolysis rate 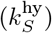 for various values of the available energy.

**Figure 3:**
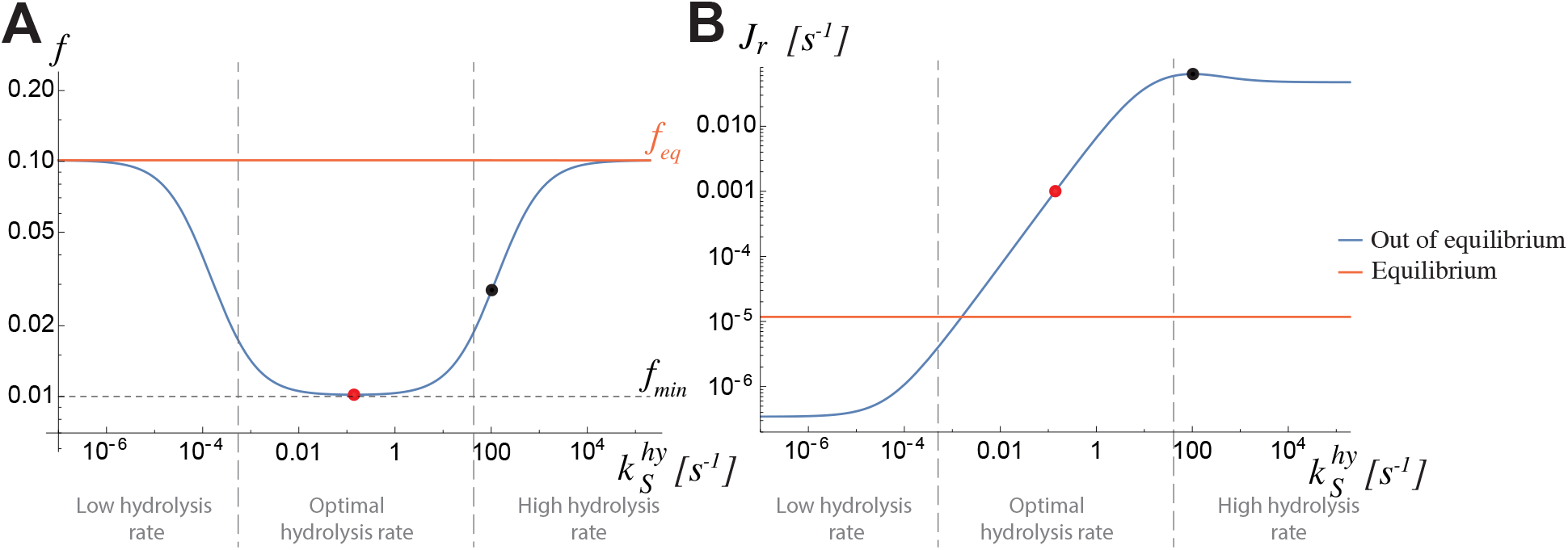
Hopfield-like selection system highly depends on available energy, *α*, and on the transition rate between *ES* and *E*S*, 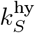. **A** Error rate and **B** production rate in function of the hydrolysis rate out of equilibrium in physiological conditions *i.e. α* = 10 (blue) and at equilibrium *α* = *α_eq_* (orange). Red dots: minimum in error rate under physiological condition (*α* = 10). Black dots: maximum in production rate under physiological condition (*α* = 10). Dashed horizontal line: theoretical limit for the error rate under Hopfield’s assumption (1).

If *α* = *α_eq_* (here *α_eq_* = 10^-6^), the error rate does not depend the hydrolysis rate, and is equal to the equilibrium predictions [3], as expected,

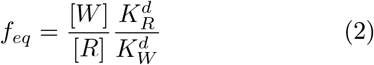

As the energy available to the system increases (increasing values of *α*), three regions can be highlighted. For very low values of the hydrolysis rate (while keeping all other rates, including exchange, fixed), the system cannot improve the error rate with respect to its equilibrium value. As a matter of fact, in this case the enzyme spends most of its time binding/unbinding the substrate while in either the *E* or *E*\* state (bound to NTP and NDP respectively) with very few hydrolysis-induced transitions between the two, as shown in Fig. 4A. Thus, no “double-checking” of the substrate takes place, *i.e.* no kinetic proofreading. As 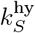 increases, there is a region where the error rate decreases, and the Hopfield result (*f* = 0.01 given the present choice of parameters) is the lower-bound that can be attained for increasing available energy. Indeed, the system is able to fully exploit the two selection steps, as highlighted in Fig. 4B. For extremely high values of the hydrolysis rate, the equilibrium error rate is again the dominant behavior, irrespective of the value of *α*. In this case, the extremely high rate of the *ES* → *E*S* transition prevents the equilibration of the *ES* complex (Fig. 4C), and thus prevents one of the two substrate-testing steps necessary for proofreading, as predicted by Hopfield [3].

**Figure 4:**
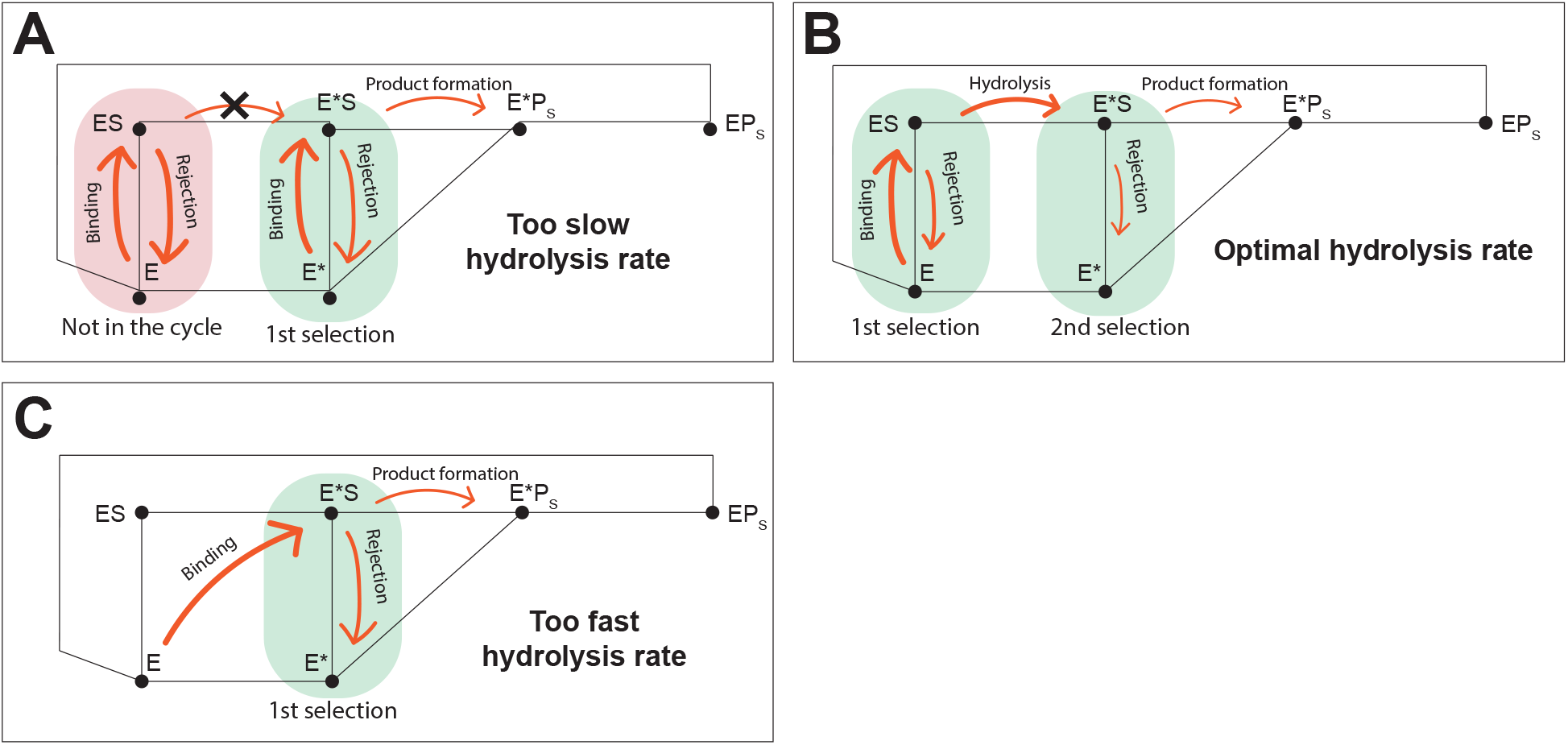
Scheme of the selection pathways following Hopfield kinetics for different values of the hydrolysis rate. **A** the slow hydrolysis rate prevent substrate from entering the selection pathway from the *E* state. Therefore the system is using only one selection step (between *E\** and *E*S,* implying a high error rate. **B** the optimal hydrolysis rate allows substrate to enter the selection pathway from the *E* state and make the most of the two selection steps leading to a low error rate. **C** with a fast hydrolysis rate, the substrate entering the selection is directly brought to the state *E* S* where it can be rejected or directly proceed to product formation. This prevents the system from benefiting from the two selection steps, leading to a higher error rate.

In addition to improving accuracy, energy consumption can also allow the process to be faster (Fig.3B). At equilibrium, *α* = *α_eq_*, the production rate is expectedly constrained to the one due to the imposed imbalance between the substrate and product concentrations. Away from equilibrium (*α* > *α_eq_*), *J_r_* can be even slower than at equilibrium for small hydrolysis rates: the increasing concentration of NTP pushes the enzyme back to *E* by nucleotide exchange, and product production is slowed. Yet, as 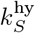 increases, hydrolysis prevails over exchange and product catalysis correspondingly increases.

Within this Hopfield-like scheme, minimizing the error rate and simultaneously maximizing the production rate is not possible at the same time. Slowing down the hydrolysis allows the system to fully exploit the two selection steps proposed by Hopfield but also slows down the passage of the substrate in the proofreading pathway. This implies a high accuracy but a production rate (red dots on Fig. 3A and B) which is not the optimal one, by orders of magnitude, in agreement with [5]. Increasing the hydrolysis rate improves the production rate up to a maximum value (black dot on Fig. 3B), at the expense of proofreading fidelity. Such trade-off between speed and accuracy, sometimes with energy dissipation, are commonly observed in information processing systems like proofreading or error correction [6, 8, 10, 12, 13], sensory adaptation [14] or polymer synthesis [15, 16, 17].

### Catalytic discrimination: Active recognition of the substrate leads to fast processes

In a kinetic proofreading scheme, accuracy is based on the repetition of energy-based discriminating steps. However, the system is forced to slow down to exploit its selection mechanism to the fullest. Indeed, the Hopfield scheme is optimal for relatively slow hydrolysis, so that the system has time to equilibrate in both *ES* and *E*S* states. The price is thus an inevitably low flux of product.

A different selection strategy could instead rely on the ability of the correct substrate to act as a catalyst for the selection process: The system would progress faster for correct substrates than for wrong ones, resulting in their **catalytic** discrimination. This selection mechanism is similar to the kinetic discrimination proposed in [5] (we here use the word catalytic instead as kinetic to avoid any confusion with the kinetic proofreading) and to the forward discrimination in [17].

This selection strategy is possible if the enzyme first recognizes the nature of the substrate (through binding) and then accelerate its uptake in the selection scheme. This approach is similar to a Maxwell Demon, whereby an appropriate action is taken only if a specific event occurs [18, 19].

In the present model, this is captured by allowing hydrolysis to be faster when the bound substrate is the correct one. At the molecular level, this scheme would require the enzyme to be able to access two different states (*e.g. ES*_1_ and *ES*_2_) that are characterized by different hydrolysis rates [20]. Substrate binding would then, allosterically, modulates the equilibrium constant between the two states in such a way that binding of the right substrate should stabilize the fast hydrolysis state more than the wrong substrate. Here, this phenomenon is coarse-grained and we are only considering the effective rate due to the inbalance between *ES*_1_ and *ES*_2_, namely 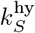, with *S* = *R* or *W*.

In the same spirit of adapting the scheme in Fig. 1 to Hopfield’s kinetic proofreading, in this case we set all the rates of the reactions involving the right and wrong substrates to be equal, apart for the hydrolysis rates (and consequently the synthesis rates) of the *ES* → *E*S* reactions. Selection is therefore only based on **catalytic discrimination** during the hydrolysis step and can thus be effective only in non-equilibrium conditions (Fig. 5). This also implies that the dissociation constants of the wrong and right substrates for the enzyme (both *E* and *E*\*) are the same and no discrimination would be possible at equilibrium (*f* = 1 if [*R*] = [*W*]).

**Figure 5:**
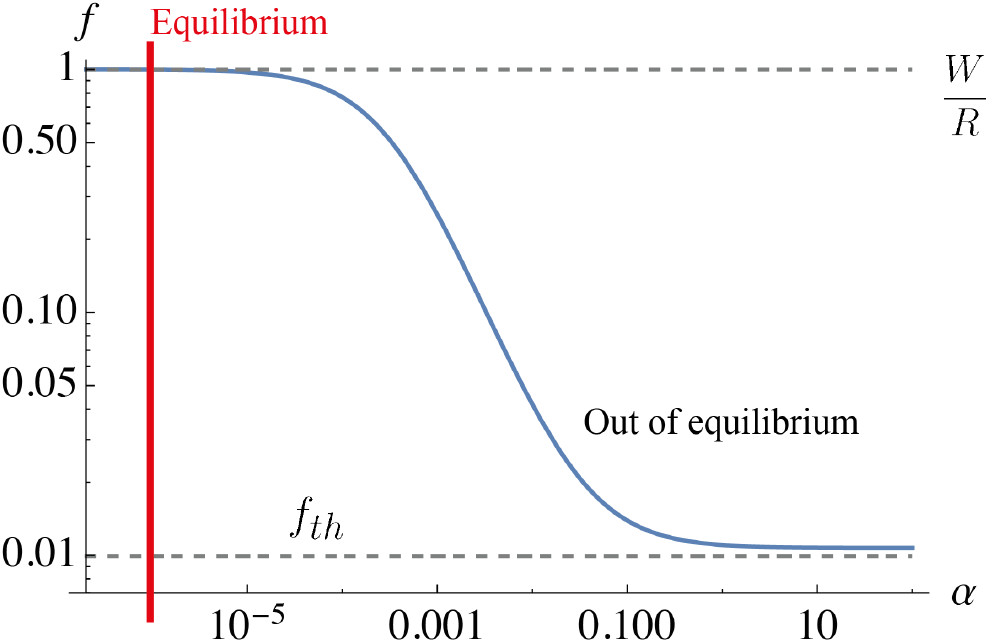
Error rate *f* in a catalytic discrimination scheme as a function of the available energy, *α*. Catalytic discrimination is only effective away from equilibrium. The upper dashed line corresponds to the error rate at equilibrium where no selection is effective, while the lower dashed line is the error rate limit (3) when the available energy is approaching ∞. The red vertical line represents the equilibrium condition with *α* = *α_eq_*.

With these assumptions and with infinite available energy (*α* → ∞ and *α_eq_* → 0), the error rate is:

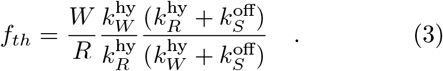

which is indeed the asymptotic limit in Fig.5 and very similar to the error rate found in [17] when considering similar discrimination mechanisms. Here 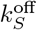 is the unbinding rate for both *W* and *R*, and 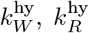, are the hydrolysis rate for *EW*, respectively *ER*. In a non-equilibrium setting, necessary to perform any kind of selection, the ability of the system to be accurate largely depends on the choice of 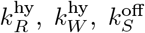 as expected from (3) and observed on Fig. 6. Clearly, when both hydrolysis rates are equal, slow or fast, no selection is possible (blue and orange line on Fig. 6A), and fast hydrolysis rates bring a higher production rate compared to slow ones. To trigger proofreading, the system needs to decrease 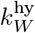 with respect to 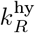, but only if the unbinding rate is set accordingly. Indeed the system starts exhibiting a significant reduction of the error rate when 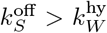 (dashed line on Fig. 6A). Overall, the minimal error rate is reached for 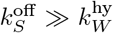. Therefore, the selectivity depends greatly on the ability to measure quickly (high 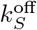) and to have an effective feedback after measurement (slower hydrolysis for wrong substrate, i.e. small 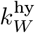).

**Figure 6:**
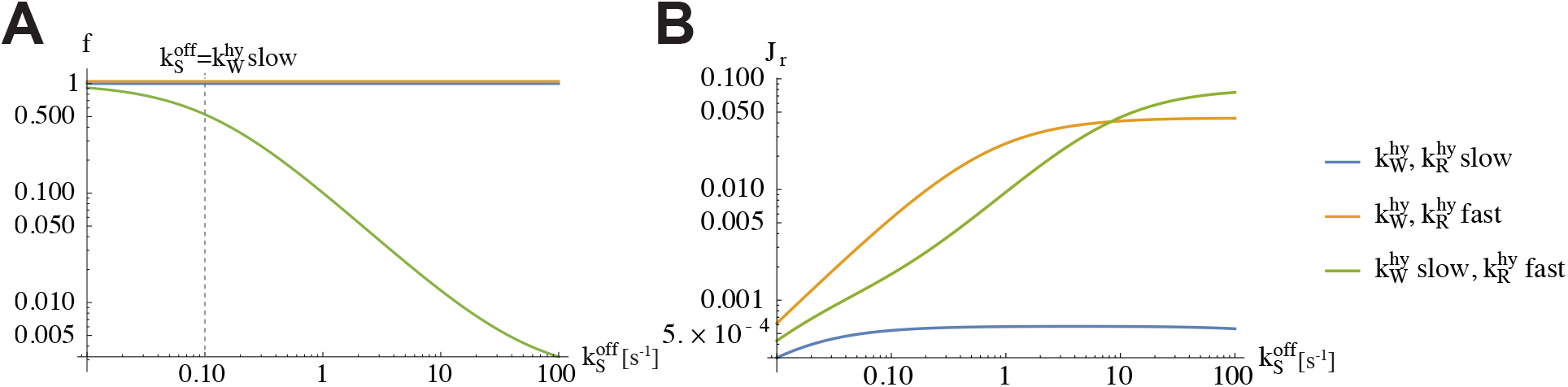
Error rate (A) and production rate (B) for slow hydrolysis rate for *W* and *R* (blue), fast hydrolysis rates for both *W* and *R* and slow hydrolysis rate for *W* and fast hydrolysis rate for *R*. For a selection mechanism based of catalytic discrimination, the fidelity is dictated by a slow hydrolysis rate for incorrect substrate, 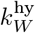, a fast hydrolysis rate for correct substrate 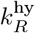, and a fast enough unbinding rate 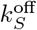.

On intuitive grounds, the system will work best when the right pathway rapidly proceeds toward the product after substrate binding, while the wrong pathway prefers to unbind the substrate before it is processed. This translates into 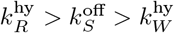.

Moreover, at variance with Hopfield’s kinetic proofreading scheme, error rate minimization could be achieved alongside maximum production rate (Fig. 6), and no compromise between accuracy and speed has to be made.

### Kinetic proofreading for accuracy, catalytic discrimination for speed

As we described above, the kinetic proofreading scheme is based on the difference in free energy between two different states. In terms of substrate binding, this implies a difference in affinity where the enzyme is more likely to be bound to the right substrate compared to the wrong one, which is discarded. To amplify this mechanism, multiple selection steps can be added separated by energy dependant steps ensuring the unidirectionality of the process. However, a downside of this strategy is the incompatibility between high production rate and low error rate. On the other hand, catalytic discrimination relies on a difference in the activation energy between forming two states of equivalent free energy. This translates into a faster process for the right substrate compared to the wrong one. This selection mechanism is only efficient in strongly nonequilibrium conditions, but without requiring a strong trade-off between speed and accuracy.

In our model, we represent the kinetic proofreading scheme as based on the difference between the unbinding rates of the right and wrong substrate with the enzyme in two different steps: before 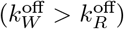 and after hydrolysis 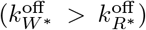. Catalytic discrimination is implemented during the hydrolysis step where we assume that the right substrate accelerates the hydrolysis compared to the wrong one. This means that the process is much faster for the right substrate than for the wrong one 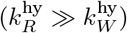.

To compare these two selection strategies, we explored the behavior of the model represented in Fig. 1 by means of two parameters that allow transitioning from one selection mechanism to the other. The first, 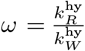, corresponds to the acceleration of the hydrolysis rate in the presence of the right substrate, 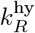, compared to the hydrolysis rate with the wrong substrate, 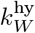, which controls the catalytic discrimination. The second, 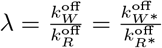, controls the kinetic proofreading *à la* Hopfield. All other parameters are completely symmetric between the two branches of the reaction network.

The region of the values of the parameters that we explored is *ω* ∈ [1,1000] and λ =∈ [1,100] (as a reference, KP indicates the selection model based on Hopfield’s kinetic proofreading scheme for *ω* = 1 and λ = 100, while C indicates catalytic discrimination for *ω* = 1000 and λ =1).

Fig. 7 reports the phase space associated to the error and production rates (*f* and *J_r_* respectively) obtained by varying *ω* and λ in three different energy conditions (*α* = *α_eq_*, 0.005,10).

**Figure 7:**
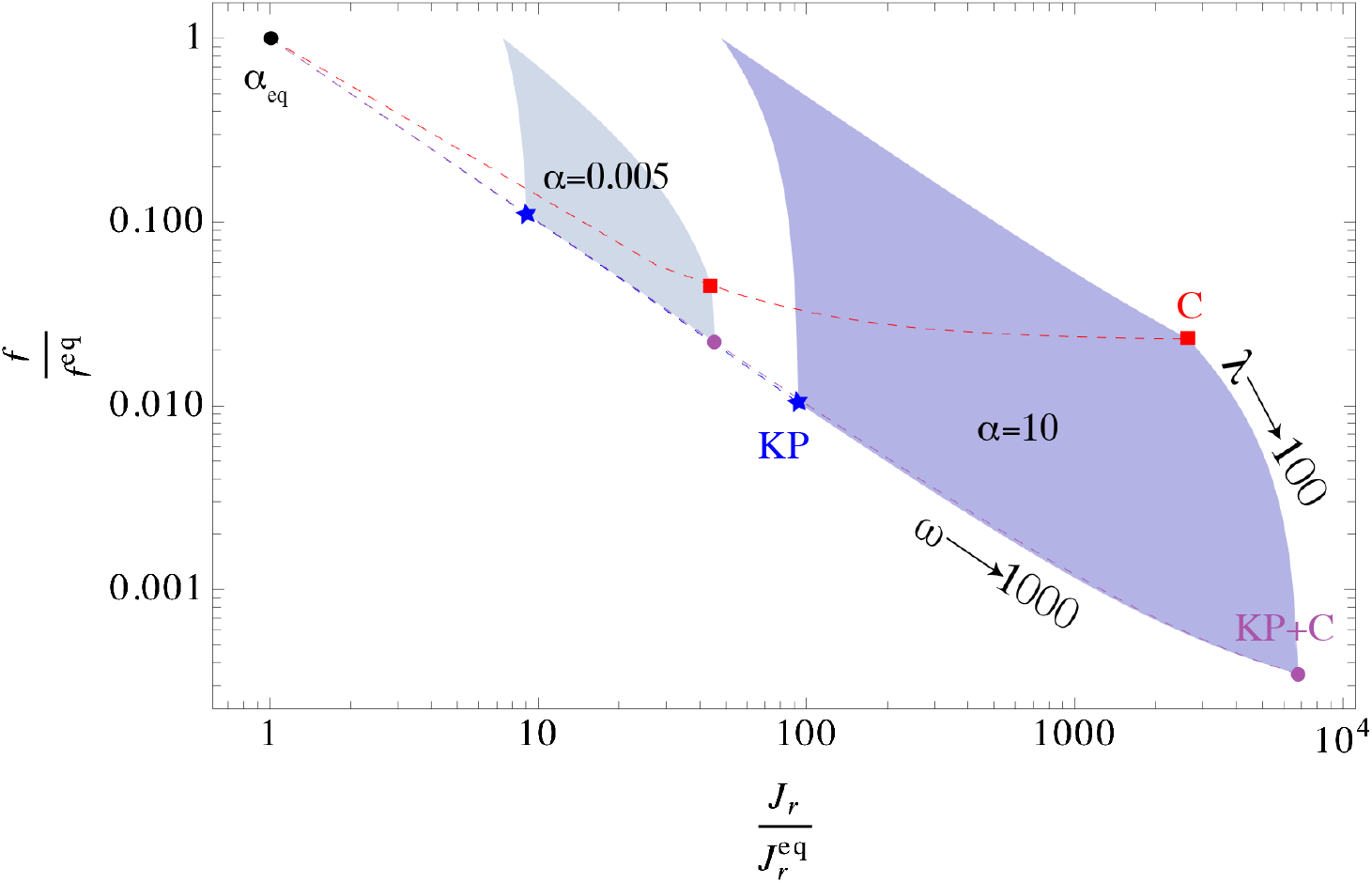
Error and production rates phase space for 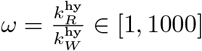, 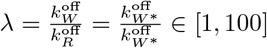 and different energy conditions (equilibrium: *α* = 10^-6^, non-equilibrium: *α* = 0.005 and *α* = 10, physiological conditions). Blue star: KP scheme (λ = 100 and *ω*=1), red square: catalytic discrimination C (λ = 1 and *ω* = 1000) and purple dot: KP + catalytic discrimination C (λ = 100, *ω* = 1000).

At equilibrium, the phase space is simply a point that captures the error rate at equilibrium, trivially related to the ratio of the dissociation constants of the right and wrong substrates. Changing the kinetic parameters cannot play any role as we saw in Fig. 5.

Increasing *α* opens a larger region of the parameter space where proofreading by both methods take place. Furthermore, it is possible to recognize the presence of a combination of KP and C (KP+C), characterized by a large difference between the hydrolysis rates (*ω* = 1000) and an important difference in the unbinding rates such that λ = 100, where the best features of both schemes coexist, namely low error rate and high production rate.

This combination of selection strategies can also be found in one of the protocols investigated in [21]. However, in their paper they only focused on obtaining the minimum error rate in particular configurations without comparing them to each other. Here we are able to show how Hopfield’s kinetic proofreading combined with catalytic discrimination is beneficial for both the accuracy and the velocity of the process.

Finally, it is worth noting the precise order of those selection mechanisms, i.e. a catalytic discrimination step in-between the two selection steps of the kinetic proofreading scheme. This order is based on biological and theoretical constraints. An energy consuming step, such as NTP hydrolysis, had to be placed in the kinetic proofreading scheme to reproduce Hopfield’s results. Furthermore, in some enzyme, the ATPase activity largely depends on substrate binding via the induced fit mechanism [22]. Hence the choice of a catalytic discrimination step during the hydrolysis. However, other combinations are possible and have been studied [5, 21].

### Selection in biological systems

The mechanisms we highlighted here are relevant to real biological systems, that crucially need both accuracy and velocity. By focusing on two of such systems, namely protein translation by ribosomes and DNA replication by polymerases, we suggest that indeed the combination of kinetic proofreading and catalytic discrimination underlies these biological processes.

#### Protein synthesis

Protein synthesis denotes the set of reactions allowing the ribosome to synthesize a protein by assembling amino acids in a precise order determined by the corresponding mRNA sequence. The ability to form a functioning protein highly depends on the ribosome accuracy in selecting the correct aminoacyl-tRNA (aa-tRNA) complex, corresponding to the targeted mRNA anti-codon. This selection is a multiple steps process.

First the ribosome binds to the ternary complex formed by aa-tRNA, the elongation factor Tu (EF-Tu) and GTP. This part is described as an initial selection step where only cognate and near cognate codons are accepted. Following the stabilization of the codon, configurational changes lead to GTPase activation allowing hydrolysis of the GTP and the release of EF-Tu. After hydrolysis, depending on the interaction between the tRNA codon and the associated anti-codon [23], the amino acid can either be added to the peptide chain or rejected. This second selection step is called proofreading.

Actually, experimental observations of the tRNA selection pathway kinetics are in agreement with the model proposed here. The ribosome is able to differentiate between cognate, near-cognate and non-cognated codons during initial selection and the proofreading step [23, 24, 25] via a two steps energy-based discrimination (kinetic proofreading), but also during GTP-hydrolysis by catalytic discrimination. Indeed, the interactions between the cognate codon and the correct anti-codon induce changes in the complex configuration which in the end lead to a fast GTPase activation followed by GTP hydrolysis [23]. This recognition, by induced-fit mechanism [24], allows to accelerate the process for cognate codon and slows it down for the near-cognate codon giving more time for the ribosome to release it. Therefore, the high accuracy is due to a discriminating process where kinetic proofreading and catalytic discrimination coexist, similar to the KP+C model.

#### DNA replication

During DNA replication, DNA polymerases are responsible for the synthesis of new DNA strands by assembling correct nucleotide bases according to the chosen DNA template.

The DNA polymerase first binds a free nucleotide (dNTP) with its polymerase domain in an open conformation. This initial binding step is depicted as a multisteps complex mechanism where accuracy of the base is controlled [26, 27]. This verification promotes conformational changes in the polymerase domain which, in turn, allows the formation of the phosphodiester bond and then the incorporation of the nucleotide. Finally, an exonuclease, or the DNA polymerase exonuclease domain, can correct mistakes by removing wrong nucleotide after their incorporation. This last step is widely regarded as a proofreading activity and was already used as a justification in the original work of Hopfield to explain the high fidelity in DNA replication [3]. Indeed, without exonuclease activity, DNA replication fidelity is decreasing from one error every 10^8^ to 10^10^ copied base-pairs [28] to one every 10^3^ to 10^6^ [29] depending on the conditions. However, this also shows that the polymerase activity is actually reaching a very high accuracy by itself. To explain this low error rate, multiple successive selection steps have been proposed. First, the initial binding step is able to reject wrong nucleotide due to a 10 to 100-fold higher affinity for the right nucleotide [30, 29]. This selection is understood to depend more on a “geometric fit” than on differences in the free energy between binding to the right and wrong nucleotide [31], but still follows the principle of the kinetic proofreading scheme. The following conformational changes leading to chemistry step of phosphodiester bond formation are promoted by an induced-fit mechanism [26], which is more that 10000 times slower for wrong nucleotide [29]. Finally, there is evidence of selection during the chemistry step [32], where the phosphodiester bond would be formed faster in presence of the correct nucleotide. While it is not yet clear if both the induced-fit mechanism and the chemistry step are able to discriminate between right and wrong nucleotide during DNA replication [26], we can consider both of them as catalytic discrimination steps, where only the initial one matters [21]. In conclusion, we once again find a strong hint that a selection mechanism relying on the same process and in the same order than our KP+C model is at work.

## Discussion

In this work, we proposed a non-equilibrium, thermodynamically consistent model of substrate selection and processing. This allows rigorously accounting for the energy which is available and which is driving the system away from equilibrium. In this framework, we tested selection based either on Hopfield’s kinetic proofreading [3] or on catalytic discrimination [5, 12], or both.

The kinetic proofreading is very effective in terms of fidelity, however the optimal fidelity comes at the price of slowing down the process by orders of magnitude. This can be compensated by the catalytic-based selection process, whose fidelity is not as high as the kinetic proofreading, but has no intrinsic velocity limitations. Including both these selection mechanisms in our model allows it to benefit from both of their advantages. Indeed, their combination allows simultaneously attaining high fidelity and fast production rate.

Given the ability of evolution to explore a vast array of possibilities, and retain the advantageous ones, it may not come as a surprise that some cellular processes exploit such a combination. Indeed, both protein translation and DNA replication, when coarse-grained to their essential selection steps, display both kinetic proofreading and catalytic discrimination steps in the same order as we suggest in our theoretical framework.

## Acknowledgments

A.M. and P.D.L.R. acknowledge support by the Swiss National Science Foundation under grant 200020_178763

## Material and Methods

### Mathematical development

#### Energy dependence and equilibrium conditions

Tuning our model out of equilibrium allows us to take advantage of the imposed directionality of reactions in the our model due to the energy consuming steps. To highlight how (and how much) the system becomes directional upon energy consumption, it is instructive to focus on two cycles from Fig. 1: *E* → *ER* → *E*R* → *E*\* → *E* and its reverse, *E* → *E*\* → *E*R* → *ER* → *E*. If the step between *ER* and *E*R* takes place over the exchange reactions (dashed double arrows in figure Fig. 1) no NTP molecule is transformed into a NDP molecule, hence no energy is consumed and the cycle must be balanced. If instead that particular step of the cycle is associated with the hydrolysis/synthesis reactions, energy consumption is at play. The unbalance between the two different ways to travel the cycle gives rise to clockwise (or anti-clockwise) directionality. This argument is made more precise using explicitly the reaction rates.

Indeed, by considering the nucleotide-less state of the enzyme (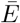 on Fig. A.1) as very short-lived and poorly populated (this approximation is useful to somewhat simplify the reaction scheme, but is not necessary for the conclusions of this work), the rates of the exchange reactions can be written as

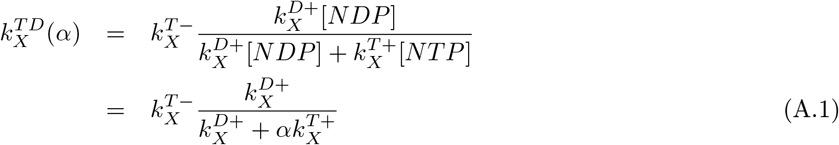

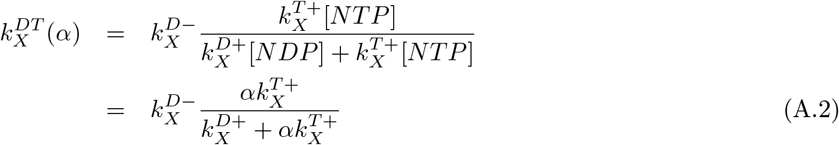

where 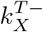 and 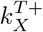 are the unbinding and binding rates of NTP, respectively (and similarly for the binding/unbinding rates of NDP), and *α* = [*NTP*]/[*NDP*]. The subscript *X* indicate the presence of a substrate (*X* = *R, W*), a product (*X* = *P_r_, P_w_*) or nothing. Their ratio is then

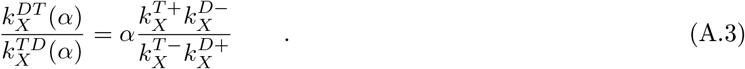

**Figure A.1:**
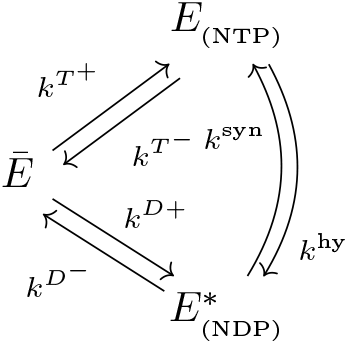
Details of the transitions between the enzyme in a NTP state (*E*) and in an NDP state (*E*\*) with an intermediate state 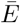 where the enzyme is not bound to any nucleotide (apo-state).

The ratio between the synthesis and hydrolysis rate can be obtained instead by considering the binding and unbinding of NTP and NDP to the enzyme, and their catalytic interconversion while bound, in an equilibrium solution of NTP and NDP: ([*NTP*]/[*NDP*])_*eq*_ = *α_eq_* (in the case of ATP and ADP, depending on the solution condition *α_eq_* ≃ 10^-8^ – 10^-6^, whereas in cellular conditions *α* ≃ 1 – 10 [11]). Applying detailed balance, it can be derived that

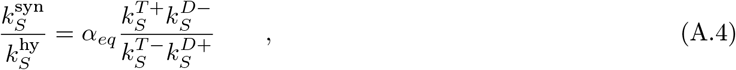

which also gives 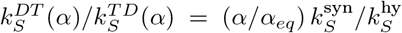. Here we are using the subscript *S* = *R, W* as the hydrolysis and synthesis only happen when the enzyme is bound to a substrate.

In equilibrium conditions (*α*/*α_eq_* = 1) the currents flowing from *ES* to *E*S* along the exchange and hydrolysis/synthesis branches are expectedly equal and opposite, so that there is no net current. Away from equilibrium (*α/α_eq_* > 1), as in cellular conditions, and if 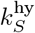 is high (as for most NTPase enzymes), the current over the hydrolysis/synthesis branch exceeds the opposite exchange current, and a net flux, which captures directionality, is then present.

#### Resolution of the model

Let *N_i_* be the *i*-th state of our system. The master equation describing the time evolution of its concentration is given by:

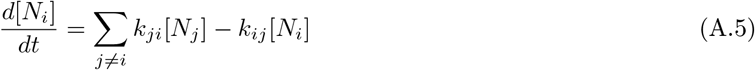

where the *k_ij_* are the transition rate from *N_i_* to *N_j_*, for all *j*.

The whole set of equations can therefore be written as

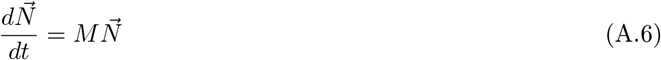

where 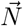 is a column vector containing the concentration of each state and *M* is the rate matrix. At steady state, the concentrations of each states are found by solving

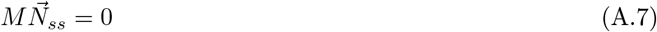

From the steady state solution, we can compute the production fluxes

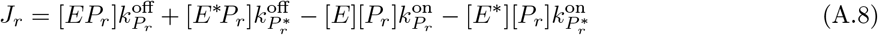

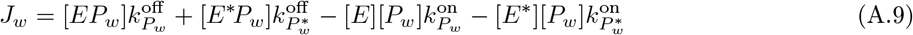

where 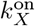 and 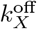 correspond to the binding and unbinding rates of the products, *X* = *P_r_* or *P_w_*. and the error rate

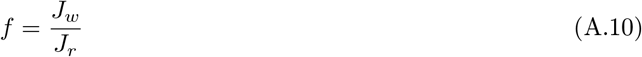

*J_r_* and *f* characterize the system accuracy and velocity, giving us a base to compare different selection methods.

### Rates for the simulations

All rates in this work were carefully chosen to respect detailed balance at equilibrium (*α* = *α_eq_*), as stated above. Therefore, among the following rates, some were freely chosen while others were constrained by the equilibrium condition.

#### Kinetic proofreading case

Numerical values to obtain results on Fig. 3.

**Table A.1:** Values of the different on and off rates used in Hopfield’s like scheme. *X* = *R,W,P_r_, P_w_* and *P_s_* = *P_r_, P_w_*.

| Conc. [$\mu\text{M}$] | | On rates [$s^{-1}\mu\text{M}^{-1}$] | | Off rates [$s^{-1}$] | | | | | |
| --- | --- | --- | --- | --- | --- | --- | --- | --- | --- |
| [R] | [W] | $k_X^{\text{on}}$ | $k_{X^*}^{\text{on}}$ | $k_R^{\text{off}}$ | $k_W^{\text{off}}$ | $k_{R^*}^{\text{off}}$ | $k_{W^*}^{\text{off}}$ | $k_{P_s}^{\text{off}}$ | $k_{P_s^*}^{\text{off}}$ |
| 1 | 1 | 95 | $10^{-3}$ | 45 | 450 | 84 | 840 | $10^{-5}$ | $10^3$ |

**Table A.2:** Rates of products formation and destruction and NTP hydrolysis and synthesis in a Hopfield’s like scheme simulation

| Production rates [$s^{-1}$] | | "Reverse production" rates [$s^{-1}$] | | Hydrolysis rate [$s^{-1}$] | Synthesis rate [$s^{-1}$] |
| --- | --- | --- | --- | --- | --- |
| $k_{RP}$ | $k_{WP}$ | $k_{PR}$ | $k_{PW}$ | $k_S^{\text{hy}}$ | $k_S^{\text{syn}}$ |
| 1 | 1 | $1.2 \times 10^{-5}$ | $1.2 \times 10^{-6}$ | varies | varies |

**Table A.3:** Rates of binding and unbinding NTP and NDP for Hopfield’s kinetic proofreading scheme

| Enzyme<br>bound to | "On rates [$s^{-1}\mu M^{-1}$]" | | "Off rates [$s^{-1}$]" | |
| --- | --- | --- | --- | --- |
| | $k^{D+}$ | $k^{T+}$ | $k^{D-}$ | $k^{T-}$ |
| ∅ | 10 | 10 | 10 | 10 |
| $S$ | 0.0018 | $5.7 \times 10^4$ | $10^{-5}$ | 0.0018 |
| $P_s$ | $2 \times 10^{-5}$ | $4.6 \times 10^6$ | $10^{-10}$ | $2 \times 10^{-5}$ |

#### Catalytic discrimination case

Numerical values to obtain results on Figs. 5 and 6.

**Table A.4:** Values of the different on and off rates used in the catalytic discrimination scheme. *X* = *R, W, P_r_, P_w_* and *P_s_* = *P_r_, P_w_*.

| Conc. [$\mu M$] | | On rates [$s^{-1}\mu M^{-1}$] | | Off rates [$s^{-1}$] | | | |
| --- | --- | --- | --- | --- | --- | --- | --- |
| [$R$] | [$W$] | $k_X^{\text{on}}$ | $k_{X^*}^{\text{on}}$ | $k_S^{\text{off}}$ | $k_{S^*}^{\text{off}}$ | $k_{P_s}^{\text{off}}$ | $k_{P_s^*}^{\text{off}}$ |
| 10 | 10 | 95 | $10^{-3}$ | varies | 84 | $10^{-5}$ | 1000 |

**Table A.5:** Rates of products formation and destruction and NTP hydrolysis and synthesis for simulation of the catalytic discrimination

| Production rates [$s^{-1}$] | | "Reverse production" rates [$s^{-1}$] | | Hydrolysis rate [$s^{-1}$] | | Synthesis rate [$s^{-1}$] | |
| --- | --- | --- | --- | --- | --- | --- | --- |
| $k_{RP}$ | $k_{WP}$ | $k_{PR}$ | $k_{PW}$ | $k_R^{hy}$ | $k_W^{hy}$ | $k_R^{syn}$ | $k_S^{syn}$ |
| 1 | 1 | $1.2 \times 10^{-4}$ | $1.2 \times 10^{-4}$ | varies | varies | $\frac{(7.98 k_R^{hy})}{k_S^{\text{off}}}$ | $\frac{(7.98 k_W^{hy})}{k_S^{\text{off}}}$ |

**Table A.6:** Rates of binding and unbinding NTP and NDP for simulation of the catalytic discrimination

| Enzyme<br>bound to | "On rates [$s^{-1}\mu M^{-1}$]" | | "Off rates [$s^{-1}$]" | |
| --- | --- | --- | --- | --- |
| | $k^{D+}$ | $k^{T+}$ | $k^{D-}$ | $k^{T-}$ |
| ∅ | 10 | 10 | 10 | 10 |
| $S$ | $5 \times 10^{-4} (k_S^{\text{off}})^{1/3}$ | $2 \times 10^{-5} (k_S^{\text{off}})^{-1/3}$ | $10^{-5}$ | $5 \times 10^{-4} (k_S^{\text{off}})^{1/3}$ |
| $P_s$ | $2 \times 10^{-5}$ | $4.6 \times 10^6$ | $10^{-10}$ | $2 \times 10^{-5}$ |

#### Comparison

Numerical values for the comparison between the two selection schemes on Fig. 7.

**Table A.7:** Values of the different on and off rates used for the comparison between kinetic proofreading and catalytic discrimination. *X* = *R, W, P_r_, P_w_* and *P_s_* = *P_r_, P_w_*. λ varies from 1 to 100, tuning the kinetic proofreading steps.

| Conc. [$\mu M$] | | On rates [$s^{-1}\mu M^{-1}$] | | Off rates [$s^{-1}$] | | | | | |
| --- | --- | --- | --- | --- | --- | --- | --- | --- | --- |
| [$R$] | [$W$] | $k_X^{\text{on}}$ | $k_{X^*}^{\text{on}}$ | $k_R^{\text{off}}$ | $k_W^{\text{off}}$ | $k_{R^*}^{\text{off}}$ | $k_{W^*}^{\text{off}}$ | $k_{P_s}^{\text{off}}$ | $k_{P_s^*}^{\text{off}}$ |
| 1 | 1 | 100 | $10^{-3}$ | 4.5 | $k_R^{\text{off}} \times \lambda$ | 8.4 | $k_{R^*}^{\text{off}} \times \lambda$ | $10^{-5}$ | 1000 |

**Table A.8:** Rates of products formation and destruction and NTP hydrolysis and synthesis, λ varies from 1 to 100, tuning the kinetic proofreading steps and *ω* varies from 1 to 1000, tuning the catalytic discrimination step.

| Production rates [$s^{-1}$] | | "Reverse production" rates [$s^{-1}$] | | Hydrolysis rate [$s^{-1}$] | | Synthesis rate [$s^{-1}$] | |
| --- | --- | --- | --- | --- | --- | --- | --- |
| $k_{RP}$ | $k_{WP}$ | $k_{PR}$ | $k_{PW}$ | $k_R^{hy}$ | $k_W^{hy}$ | $k_R^{syn}$ | $k_W^{syn}$ |
| 100 | 100 | 0.012 | $0.012/\lambda$ | $k_W^{hy} \times \omega$ | 0.1 | $0.19 k_R^{hy}$ | 0.019 |

**Table A.9:**
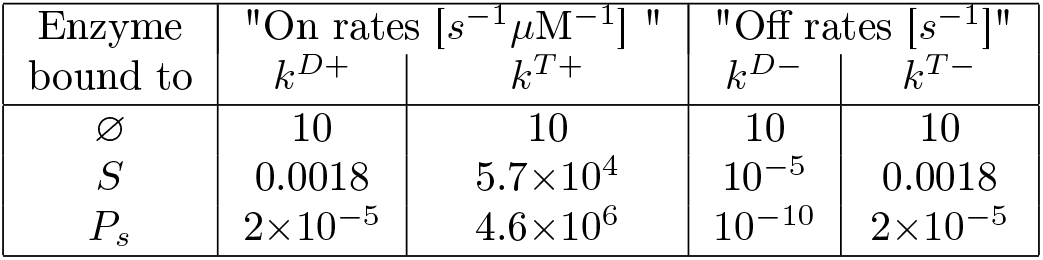
Rates of binding and unbinding NTP and NDP for comparison between kinetic proofreading and catalytic discrimination

